# Morphological estimation of Cellularity on Neo-adjuvant treated breast cancer histological images

**DOI:** 10.1101/2020.04.01.020719

**Authors:** Mauricio Alberto Ortega-Ruiz, Cefa Karabağ, Victor García Garduño, Constantino Carlos Reyes-Aldasoro

**Affiliations:** Universidad del Valle de Mexico, Campus Coyoacán C.P. 04910 CDMX Mexico; School of Mathematics, Computer Science and Engineering City, University of London, London EC1V 0HB, UK; Universidad Nacional Autonoma de Mexico, Mexico

## Abstract

This paper describes a methodology that extracts morphological features from histological breast cancer images stained for Hematoxilyn and Eosin (H&E). Cellularity was estimated and the correlation between features and the residual tumour size cellularity after a Neo-Adjuvant treatment (NAT) was examined. Images from whole slide imaging (WSI) were processed automatically with traditional computer vision methods to extract twenty two morphological parameters from the nuclei, epithelial region and the global image. The methodology was applied to a set of images from breast cancer under NAT. The data came from the BreastPathQ Cancer Cellularity Challenge 2019, and consisted of 2579 patches of 255×255 pixels of H&E histopatological samples from NAT treatment patients. The methodology automatically implements colour separation, segmentation and morphological analysis using traditional algorithms (K-means grouping, watershed segmentation, Otsu’s binarisation). Linear regression methods were applied to determine strongest correlation between the parameters and the cancer cellularity. The morphological parameters showed correlation with the residual tumour cancer cellularity. The strongest correlations corresponded to the stroma concentration value (r = −0.9786) and value from HSV image colour space (r = −0.9728), both from a global image parameters.

## 1 Background

Digital pathology has recently become a major player in cancer research, disease detection, classification and even in outcome prognosis (1; 2). Perhaps the most common imaging technique is Hematoxylin and Eosin (H&E), where H stains nuclei blue and E stains cytoplasm pink. Also, Immunohistochemistry methods (IHC) that use antibodies to stain specific antigens or proteins in the tissue are more specific and can complement H&E. With the spread use of whole slide digital scanners (3) numerous of histopathology images have become available for research purposes. Computer Aided Diagnosis (CAD) has become an important research area (2) and is based on the quantitative analysis to grade the level of the disease, done by processing the image with both, computer vision algorithms and deep-learning techniques, that performs separation, segmentation, counting and classification of positive cells of cancer disease (4). Quantitative information is being also used to explore new clinical-pathological relationship with the data (5). For example, a better understanding of mechanisms of the disease evolution process (1), as the image contains high amount of information at a cell level, fundamental prognosis information is contained in pathology image data (6). Also there have been attempts to correlate morphological parameters of cancer images to a specific clinical behaviour (5; 7). This is the main objective of present paper.

Among all cancer diseases in women, Breast Cancer has the most common new occurrences. For that reason, different analysis imaging methods are employed to study this type of cancer images (8). In 2018, 30% of new cases in the US reported in women were breast cancer and also represents the second place death with 14% cases (9). Common treatments for breast cancer are surgery, radiation, chemotherapy or targeted therapy. Breast Cancer Neo-adjuvant treatment (NAT) is a therapy for advanced disease cases that gives useful information for prognosis and disease outcome. Response is evaluated by comparing tissue samples before, during and after treatment. This is done by physical examination, ultrasonography and mammography. An accuracy estimation of residual cancer is needed to asses therapy response, mainly based on tumour size. The main advantages of the use of NAT are reduction of tumour size and also to determine a prognostic indicator of final therapy (10).

Cellularity is defined as the fraction of malignant cells within tumour bed and is calculated as the rate of the total malignant area by the total image area, i.e. the percentage area occupied by malignant cells per sq. mm. It has been reported that cellularity is an accurate indicator of efficacy of treatment (11; 12). Cellularity is calculated manually by pathologists, based on visual analysis of nuclei cell within the tissue micro-environment which is a time consuming task, for that reason there is a motivation to develop efficient methods to estimate cellularity based on the use of image processing and artificial intelligence (AI) algorithms (10).

The motivation of this paper is to present an automated computer methodology to process series of WSI from Breast cancer NAT images and correlate the morphological parameters obtained with cellularity of residual tumour. This is a first step in the development of an automated cellularity computation system.

## 2 The Data

The data set for this study was obtained from the Breast SPIE challenge 2019 and was collected at the Sunnybrook Health Sciences Centre, Toronto. It comprises a set of WSI which have been stained with H&E (10)

Images were extracted from patients with residual invasive breast cancer on re-section specimens following neo-adjuvant therapy. The specimens were handled according to routine clinical protocols and WSIs were scanned at 20× magnification (0.5*µ* m/pixel).

In the data set, there is a training set of 2579 patches of size 255 ×255 extracted from the above WSIs which has a reference standard from 2 pathologists that indicate residual cellularity and is considered as the reference Ground Truth (GT) of this study. Each patch has a score on a scale from 0 to 1. Zero means no cancer cells are present and one indicate the image is completely full of malignant cells according to cellularity computation. Bottom rows in Figure 1 present samples of images from cancer cellularity zero to one.

**Fig. 1.**
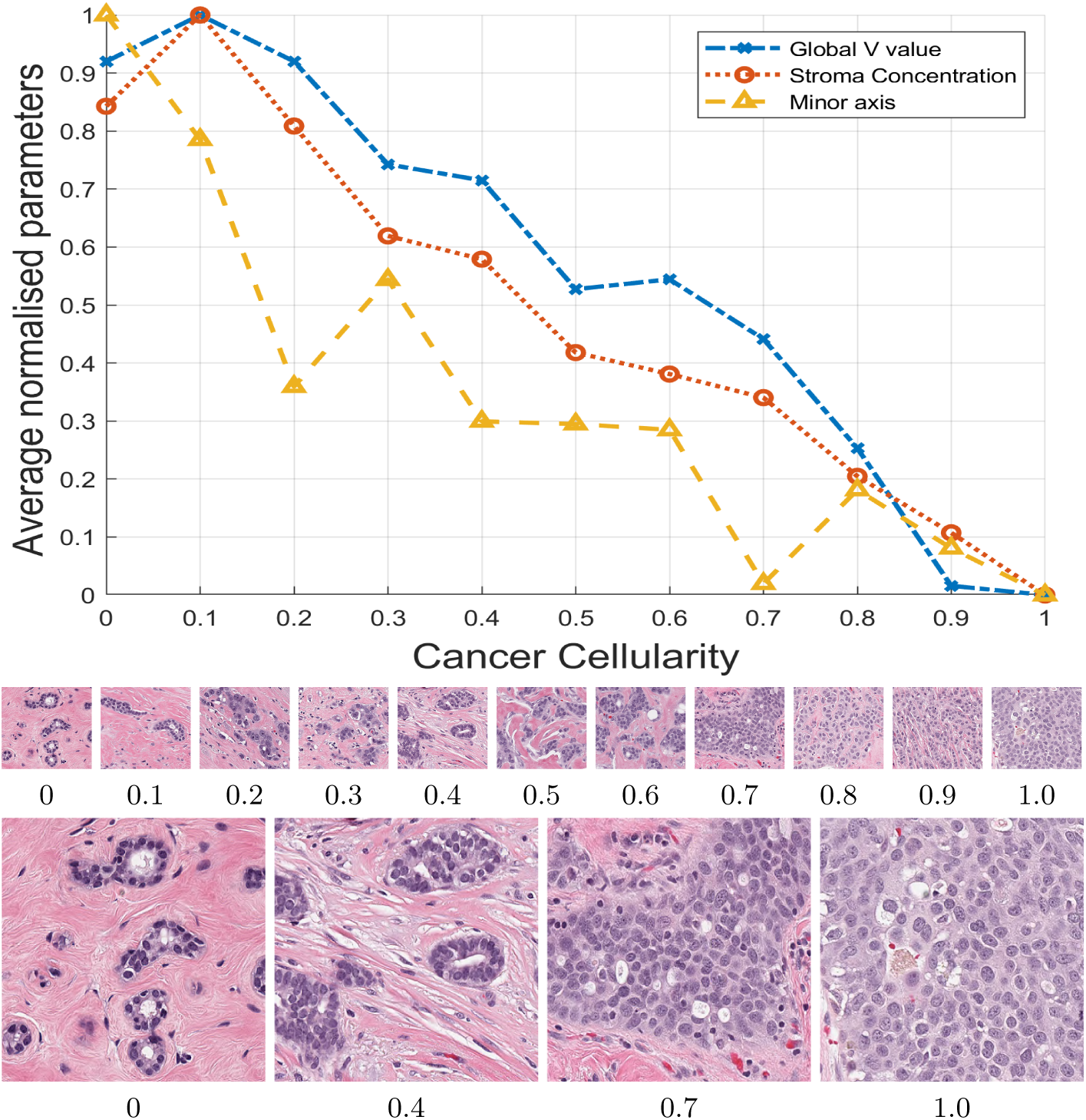
Graphical display of the main morphological parameters against the cellularity of the images. Cellularity was manually ranked by a pathologist and the parameters automatically extracted. Plots represent average parameter value from the whole 2579 processed images. All the parameters are normalised with maximum value to be compared in the same graph. Lines correspond to 3 of the parameters with strongest correlation: Global stroma filtered region (o), minor axis(^) and Value concentration in region (+). Figure indicates a relationship between respective parameter and cellularity. Thumbnails show images with cellularity values from zero to one and magnified versions of cases with values 0, 0.4, 0.7 and 1.

## 3 Description of the method

The methodology processed automatically the 2579 images using three regions of interest: inner segmented cell region, neighbourhood around segmented cell and the full image. A master control routine is responsible of selecting one by one the corresponding image to be processed and also saves the result in the final data file. The methodology allows two operational modes, a supervised and unsupervised mode. Supervised mode is configured to analyse segmented cells manually, useful to classify cells into different categories and generate data to train machine learning methods. Unsupervised mode processes the full set automatically and generates different files with all image cell parameters. All the coding was was implemented in Matlab® (The Mathworks™, Natick, USA) 2019b version, with functions from the digital image processing, statistical and machine learning toolboxes.

In the next step, digital imaging and computer vision algorithms are implemented to process the image at the three processing regions (inner nuclei, regional and global level). When all the processing is concluded, a large amount of morphological parameters were obtained and those corresponding to Celullarity values equal to (0, 0.1, 0.2 …1.0) were selected. Their mean, standard deviation, maximum, minimum values were computed, and a linear regression analysis was determined to estimate strongest correlation to cellularity. From all the parameters analysed the 22 with strongest correlation are presented in Table 1. These parameters are described according to the processing region. First, the morphological parameters related to segmented nuclei, are computed with image processing of binary black and white image object: eccentricity, roundness, major axis, minor axis and perimeter. Then, parameters computed at neighborhood region in its original RGB image and in transformed HSV colour map image. These parameters are: nuclei density concentration (as explained later in level processing region), Hue, Value, and Saturation histograms of the regional HSV colour map image. Finally, parameters computed from the full image are: HSV average values from HSV components, nuclei concentration, basement concentration and stroma average concentration determined at output of a special pink filter implemented.

**Table 1.**
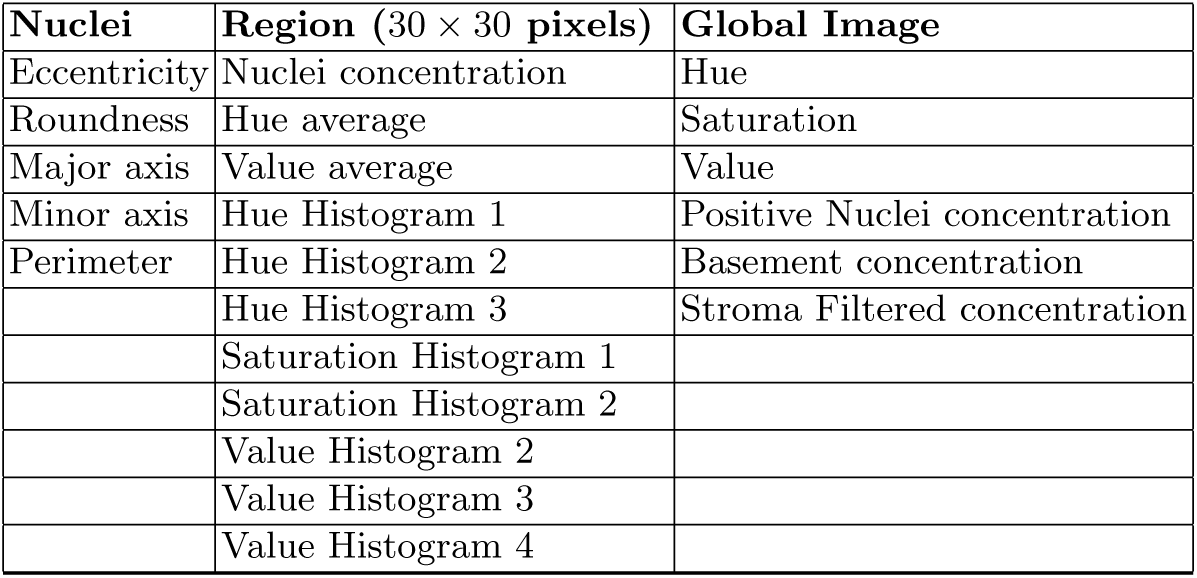
Morphological parameters with which Cancer cellularity had the strongest correlation. Column division corresponds to the processing region: nuclei, regional and whole image.

| Nuclei | Region (30 × 30 pixels) | Global Image |
| --- | --- | --- |
| Eccentricity | Nuclei concentration | Hue |
| Roundness | Hue average | Saturation |
| Major axis | Value average | Value |
| Minor axis | Hue Histogram 1 | Positive Nuclei concentration |
| Perimeter | Hue Histogram 2 | Basement concentration |
|  | Hue Histogram 3 | Stroma Filtered concentration |
|  | Saturation Histogram 1 |  |
|  | Saturation Histogram 2 |  |
|  | Value Histogram 2 |  |
|  | Value Histogram 3 |  |
|  | Value Histogram 4 |  |

The correlation of some of the morphological parameters with cellularity are illustrated in Figure 1. A series of images with different levels of cellularity are illustrated below the graph. The processing methods implemented at each region are described in the following section.

### 3.1 Processing methods at nuclei level

Once the corresponding image is selected, colour separation, nuclei segmentation and binarisation (1) were implemented. The image is first converted to HSV colour space and then by K-means clustering (13) three main colour images are separated. The detected images are *Ip* (for pink), *Ib* (for blue) and *Iba* is the remaining background colour component. Then, the selected positive nuclei image *Ib* is binarised. Different segmenting methods have been reported in the literature (14) and a segmenting method based on two simple steps is proposed: first cell regions are enhanced and background region is weakened, which is obtained by means of equation 1. Next, the enhanced image *In* is transformed to a binary image by thresholding with Otsu’s algorithm (15). Binarisation threshold was selected adaptively in dependency of strongest correlated parameter with cellularity, obtained after a first computation trial of this methodology. Enhancement procedure is as follows: The RGB image is converted to gray level image, say *Io*, then a median filter was applied to get image *Im*, and the enhanced regional image *In* is obtained by equation

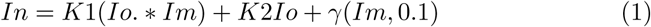

where *γ* is the gamma correction of median filter image *Im* of parameter 0.1. K1 and K2 are constants (0, 1). K1 enhances cellular regions, K2 reduces original gray level value and low gamma function weakens full image to reduce background intensity. Figure 2 shows an example of enhanced image and segmented binary result. A visual inspection of this segmentation procedure compared to binarisation of original gray image indicates a better cell sementation when applied to the full 2579 patch set. Binarisation was complemented with watershed segmentation (16) to separate touching cell elements. The binary image is then analysed by traditional morphological algorithms to obtain its main parameters: area, perimeter, roundness, eccentricity, centroid X, centroid Y, major axis, minor axis and orientation angle. A mean area size was determined and objects lower than 15% mean size were discarded, this value can be adjusted by the software user. Also a bounding box coordinates of segmented cell body was determined and inner cell parameters were determined: texture and mean HSV values of the body.

**Fig. 2.**
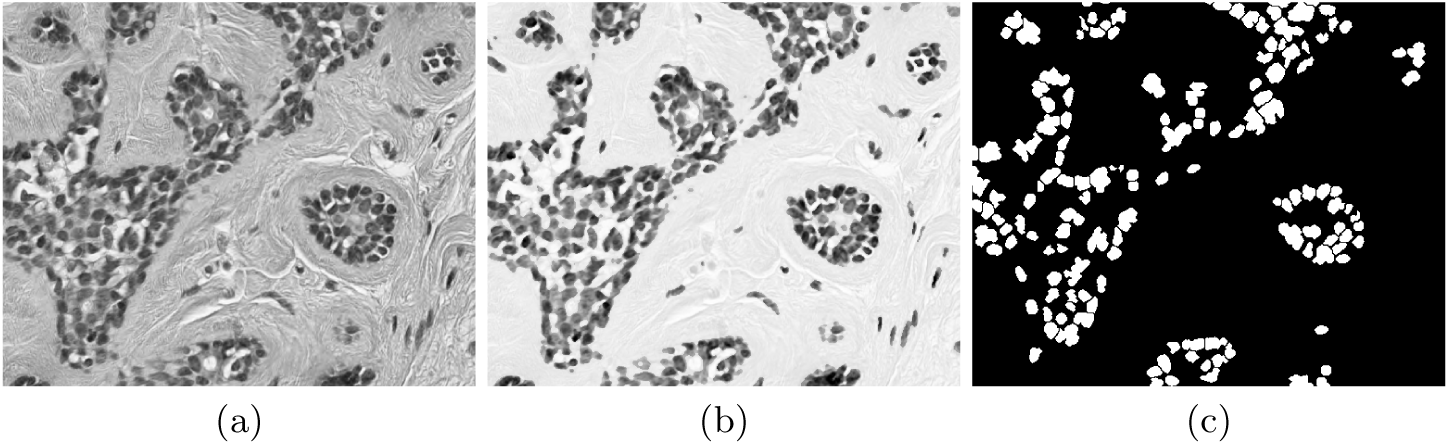
Enhancement of nuclei area. (a) Original gray level image. (b) Enhanced image, notice that the nuclei region is darker whilst background becomes brighter. (c) Binary image obtained with an adaptive threshold value estimated directly from strongest correlated parameter.

### 3.2 Regional level processing

Characteristics of the neighbourhood parameters surrounding the segmented cell were calculated at different window sizes around the segmented nuclei: 30× 30, 70× 70, 90 ×90 and 120 ×120 pixels. The parameters estimated are the corresponding regional concentration of *Ip, Iba, Ib* and HSV calculated from the binary image. This concentration is determined by the ratio of total white pixels *T*_*W*_ by total pixels *T*_*P*_ of window neighborhood region.

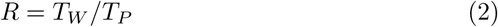

Figure 3 shows examples of regional density computation, red and brown areas in figure (d) and (g) indicate highest concentration whereas white areas indicate low concentration. Each colour square is a 30 × 30 window sub-image. Two images are presented : *Ip* and *Ib*. The regional analysis included also four bins histogram into the window sub-image region from Hue, Saturation and Value window components.

**Fig. 3.**
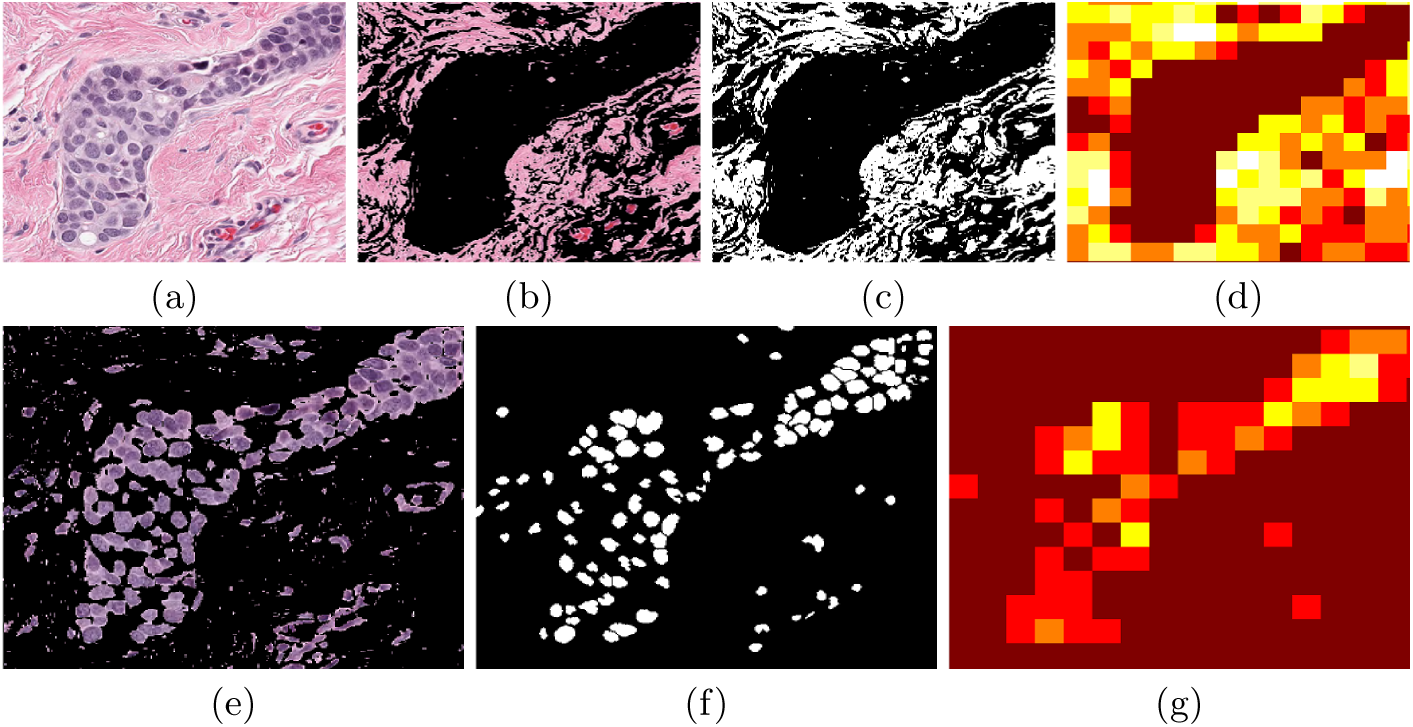
Regional analysis of the data. (a) Original image. (b) *Ip*, stroma image obtained from colour separation. (c) Binarised *Ip* image. (d) Regional image concentration in which white colour indicates the lowest and dark brown is the highest region concentration. (e) *Ib* nuclei image obtained from colour separation. (f) Binarised *Ib* image. (g) Regional density concentration. This example was processed at 30 × 30 pixels window.

Some healthy tissue images (with low cellularity), show vessels which form a defined ordered region. This is formed of positive cells grouped into the same epithelial tissue, that can be seen as nuclei dark blue within epithelial of lower blue colour intensity as seen in Figure 4. These regions are detected by a method based on simple Set theory. Let U be the full image, D be a cluster of cells (vessel), say D ⊆U ≠ ∅. Then D regions are obtained by subtracting background image *Iba* obtained after colour separation from original image U. Morphological parameters from every cluster are estimated: total cluster area, cluster roundness, total number of cells inside cluster and distance from cells centroid to its corresponding cluster centroid.

**Fig. 4.**
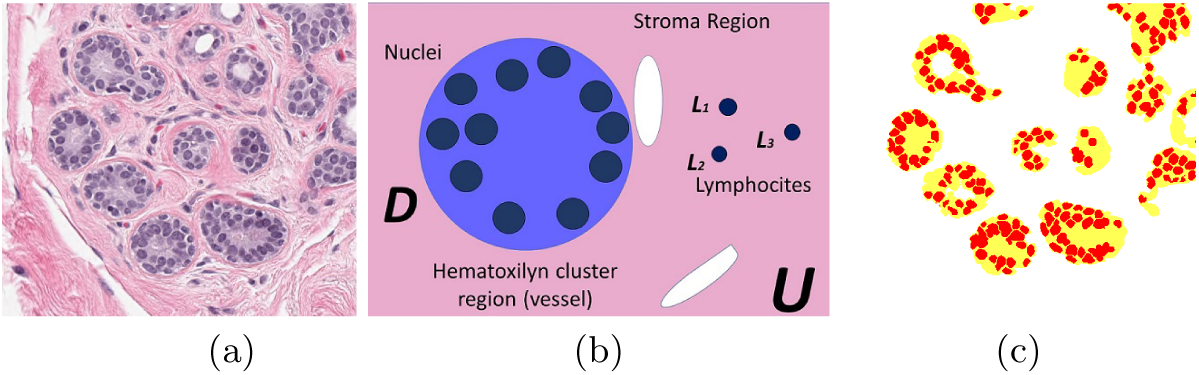
Example of vessels defined as a cluster region. (a) Original image. (b) Graphical description of a cluster region. (c) Clusters detected by the methodology labelled in different colours.

### 3.3 Global image processing

Parameters from the complete full image were also computed using original colour image and transformed HSV colour space images. The average value of full image component were calculated, then all of the cells from the full image have the same value assigned. Similarly from the separated blue, pink and background colour the average value were determined. Also, a pink colour filter was implemented by means of the Matlab®(The Mathworks™, Natick, USA) colour thresholder application which extracts stroma region from the image and average value is computed.

## 4 Results

The full set of 2579 images were processed by the methodology here described. The 22 parameters from every single segmented cell were estimated and all average values calculated and stored. Images corresponding to Cellularity values equal to (0, 0.1, 0.2…0.9, 1.0) were extracted and a matrix of the corresponding value vs. cellularity was computed. Correlation between every parameter and cellularity by linear regression analysis was computed and a summary with strongest correlation parameters is presented. Three regions of interest were processed: inner segmented cell region, regional neighbourhood around segmented cell and the full image. Figure 5 shows samples of parameters statistical behaviour with strong correlation with cancer Cellularity after analysing thousands from each processed region. Morphological information from segmented nuclei region showed strongest correlation values corresponds to: minor axis (r = 0.8939), perimeter (r = 0.8786), eccentricity (r =− 0.8771) and area (r = 0.8462).

**Fig. 5.**
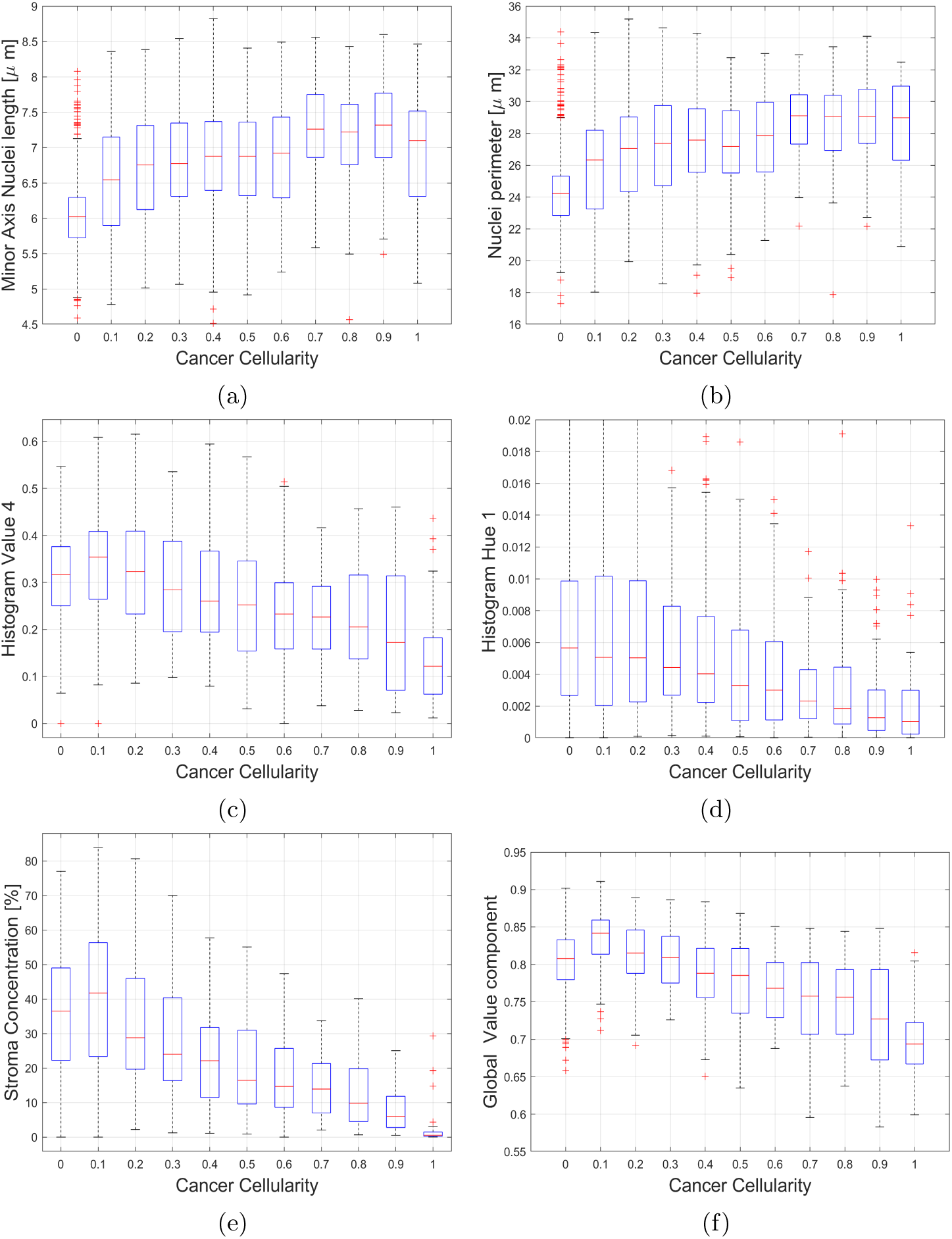
Box plots of selected parameters with strongest correlation from each ROI. From inner segmented cell region are (a) Minor axis and (b) Nuclei Perimeter. From neighbourhood around segmented cell (c) Value and (d) Hue concentration. From full image (e) Stroma filtered average value and (f) Value concentration component from HSV colour map image.

From neighbourhood region around segmented cells, concentrations of stroma, nuclei, epithelium and basement tissue were computed. Positive nuclei concentration surrounding the cell indicate the strongest correlation (r = 0.0.8119). Also RGB image section was transformed to HSV colour map image to determine HSV components by means of four bins histograms. Concentrations of Hue (r = −0.8454), Value (r = −0.8734) and most of HSV histograms indicate strong correlation. Hue histogram have strong correlation in all its 4 bins values histograms (r1 = −0.9659, r2 = −0.8997, r3 = 0.8684, r4 = −0.86), and only first two bins of Saturation and last 2 bins of Value histogram. From the cluster (vessels) region no relevant correlation was found, also a supervision inspection of selected clusters by the methodology showed that both images with high cellularity having malignant cells present random cluster regions which affect this parameter result.

The full RGB image was transformed to HSV colour map and global values of Hue (r = −0.9068), Saturation (r = 0.9) and Value (r = −0.9728) (r = −0.9068, r = 0.9, r = −0.9728) have strong correlation as well as basement concentration (r = −0.8192). Finally, stroma concentration of the full image obtained from a pink colour filter showed the strongest correlation of all the parameters under study (r = −0.9786). For this reason, this parameter was selected as a reference to determine the adaptive threshold value of positive cells binary segmentation.

## 5 Conclusions

A computer methodology that processes automatically H&E histopathology digital images to extract main morphological parameters at a cell, regional and global level is presented in this paper. The methodology was tested to process breast cancer images under neo-adjuvant treatment and the results indicate 22 key morphological parameters are strongly correlated with cellularity. These were used to train machine learning algorithms (17) to implement a system for automated cellularity estimation, with satisfactory results submitted at the challenge contest. The strongest related parameter was stroma density, as reported by Beck et al. (7) the histology of stroma correlates with prognostic in breast cancer. This result suggest the importance of a deeper image analysis of this region.

